# Bayesian Structural Time Series for Biomedical Sensor Data: A Flexible Modeling Framework for Evaluating Interventions

**DOI:** 10.1101/2020.03.02.973677

**Authors:** Jason Liu, Daniel J. Spakowicz, Garrett I. Ash, Rebecca Hoyd, Andrew Zhang, Shaoke Lou, Donghoon Lee, Jing Zhang, Carolyn Presley, Ann Greene, Matthew Stults-Kolehmainen, Laura Nally, Julien S. Baker, Lisa M. Fucito, Stuart A. Weinzimer, Andrew V Papachristos, Mark Gerstein

## Abstract

The development of mobile-health technology has the potential to revolutionize personalized medicine. Biomedical sensors (e.g. wearables) can assist with determining treatment plans for individuals, provide quantitative information to healthcare providers, and give objective measurements of health, leading to the goal of precise phenotypic correlates for genotypes. Even though treatments and interventions are becoming more specific and datasets more abundant, measuring the causal impact of health interventions requires careful considerations of complex covariate structures as well as knowledge of the temporal and spatial properties of the data. Thus, biomedical sensor data need to make use of specialized statistical models. Here, we show how the Bayesian structural time series framework, widely used in economics, can be applied to these data. We further show how this framework corrects for covariates to provide accurate assessments of interventions. Furthermore, it allows for a time-dependent confidence interval of impact, which is useful for considering individualized assessments of intervention efficacy. We provide a customized biomedical adaptor tool around a specific Google implementation of the Bayesian structural time series framework that uniformly processes, prepares, and registers diverse biomedical data. We apply the resulting software implementation to a structured set of examples in biomedicine to showcase the ability of the framework to evaluate interventions with varying levels of data richness and covariate complexity. In particular, we show how the framework is able to evaluate an exercise intervention’s effect on stabilizing blood glucose in a diabetes dataset. We also provide a future-anticipating illustration from a behavioral dataset showcasing how the framework integrates complex spatial covariates. Overall, we show the robustness of the Bayesian structural time series framework when applied to biomedical sensor data, highlighting its increasing value for current and future datasets.

## INTRODUCTION

### Background

The wearable sensor market was valued at $10.8 billion USD in 2019 and is expected to triple in value over the next five years (1). Investment is bolstered by the great potential for advancing personalized medicine in the near future (2–4). Whereas medicine has previously focused on determining the right interventions, it is now more focused on for whom and when (5). Identifying the right time for and timing of treatments remains relatively understudied, but this trend is expected to change soon as large streams of sensor data are released (6). As sensor technology develops, data-rich features such as physical, chemical, behavioral, and biological variables will be measurable. In addition to time series data, spatial information is becoming more popular as well (7), all of which can be used for a more detailed understanding of sensors. A survey of distinct types of data are presented in Figure 1.

**Figure 1.**
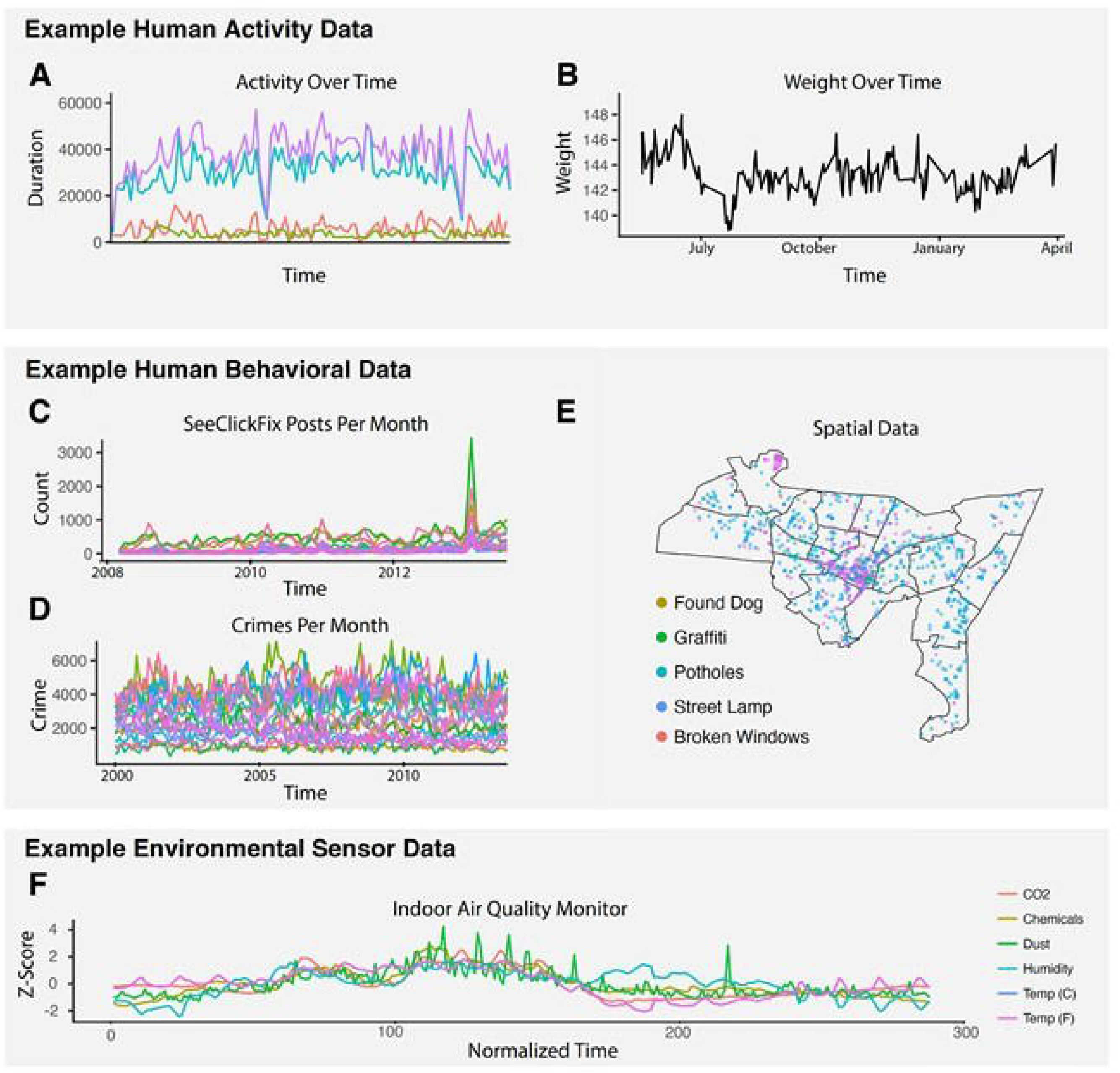
A) MOVES data set showing different exercise activities overtime, with each color representing a different activity. B) Weight overtime from the Withings data set. C) An example ofhuman behavioral data, the use ofSeeClickFix, over time. D) Various neighborhoods in New Haven and their crime data over time. E) Spatial map ofNew Haven illustrating the use of SeeClickFix and the associated geographical data. F) Various measurements taken from an indoor air quality sensor averaged over a day across a month.

Though increasing amounts of sensor data is being released, there still is a paucity of analytical methods in the biomedical field that can accommodate the complex covariate structures as well as the temporal and local trend considerations necessary to analyze these longitudinal data. Existing methods applied to these situations are limited. They address longitudinally observed patterns that could inform the timing of an intervention but do not evaluate the impact and efficacy of such an intervention.

While the nature of data from sensors and wearables will vary depending on the context, most of them share certain properties. A variety of such data can be seen in Figure 1, all of which come from various contexts. It is important to discern these properties and to establish a flexible model for emerging sensor and wearable data, as it will have broad implications in fields such as personalized medicine (2). Therefore, in this study we aim to contextualize a model and establish a flexible statistical framework to model various sensor and mobile health data. The model we adapted here is a combination of the Bayesian structural time series model and the Causal Impact model from Google (8). The principles of this modeling framework stem from Bayesian inference and the analysis of time series data, which have been well established for decades (9). While the idea of using Bayesian inference models has been extensively used in fields such as finance (10), ad campaigns (11), and marketing (12), they have rarely been used in the context of wearable and sensor data analysis to specifically evaluate impact intervention over various time periods. A survey of the application of Bayesian modeling of biosensor time series data yields few results. Therefore, we hope to apply this widely accepted statistical framework to the specialized context of biomedicine. By doing so we showcase the effectiveness when applied to a variety of sensor data we collected and accurately assess the impact that various interventions have on individuals.

### Modelling Framework

To establish an intuitive modeling framework for data taken from sensors and other mobile health sources, it is important to lay out the structure of how the framework is formed as well as any assumptions that may be essential for the model. First, we must recognize that the dataset is a time series data with a known response variable that evolves over time. For example, this could be a person’s weight. This response variable could be a direct measurement taken from a sensor, or it could be a derived value – some scalar calculated from various underlying variables measured by a device – such as calories burned (13) or a Nike Fuel Score (14). The response variable’s evolution in time is important, as it should be dynamic and change based on covariates and interventions. In the example of weight, the measured weight fluctuates over time based on things such as seasonality (15), temperature (16), or diet (17).

Because we are interested in the impact that an intervention has on such a variable, an important assumption is that we know when the intervention occurs as well as the duration. There exist methods that detect intervention times (18,19) though to jointly assess the impact would result in a significant statistical disadvantage due to the multiple hypothesis corrections since the predictions from these models only account for one time point rather than a set of time points. Therefore, by having prior knowledge on the time of occurrence and duration of the intervention, we are able to more accurately assess the impact on an individualized level.

Given a specific intervention and its duration, we can split our time series into two segments – the pre- and post- intervention periods. These periods are used to describe all variables, known and unknown, as well as any parameters associated with them. For the example of weight and diet, the pre-intervention would be the period before the diet starts and the post-intervention is the period after the diet starts. This is distinct from “post-cessation of intervention” (e.g., return to unhealthy eating following several weeks of dieting) which is not addressed by our present model.

Our goal is to model the pre-intervention period using covariates to most accurately assess any changes in behavior in the response variable before any intervention has occurred. We also assume that any covariates used in our model should not be affected by the intervention, to provide a similar control in the post-intervention period. Though the post-intervention covariates themselves may change in value, the assumption is that they are derived from the same distribution as in the pre-intervention. Finally, using the model derived from the pre-intervention period and the covariates in the post-intervention, we can calculate a counterfactual in the post-intervention. The counterfactual predicts the response variable without intervention. It serves as a baseline to compare the actual observation in the post-intervention and ultimately is what we use to calculate the impact of an intervention. Compared to linear models, this framework allows for an evolving measure of impact, due to the dynamic confidence interval for the difference between counterfactual and observation that is inherent to using Bayesian structural time series. This temporal consideration, in addition to the more common advantage of using hyper-parameters and priors, is an important consideration for the Bayesian framework, and sets it apart from other models.

This Bayesian structural time series framework can make use of complex covariate structures, which is useful and necessary to get an unbiased measure of impact. Two main types of covariates can be used: those that have known effects on the response and those that can account for hidden effects. The first type refers to covariates that could be correlated to the response, or cause changes to the response unrelated to the intervention. In relation to our example about weight, covariates of this type may include temperature, weather, or season. Since it is very unlikely to know all covariates of the first type that may perturb the response variable, we also introduce the notion of a second type of covariate, termed a “paired covariate”. The paired covariate is an independent stream of sensor data of the same type as our original data in question. They could be collected in different locations, but the ideal case is they receive little to no treatment or intervention. If we imagine a scenario where the roommate of our subject is subject to many of the same external factors, but the roommate does not participate in the dieting, then this roommate’s weight could be considered the paired covariate. Since both the original data and paired covariate share underlying biases, by using the paired covariates we can determine the true impact of the intervention.

As a whole, the Bayesian structural time series framework addresses many of the limitations of other methods and serves to advance the field by providing a structured method for analyzing data from sensors and mobile health sources. We give more details in the Statistical Formalism section and provide a schematic of the framework in Figure 2.

**Figure 2.**
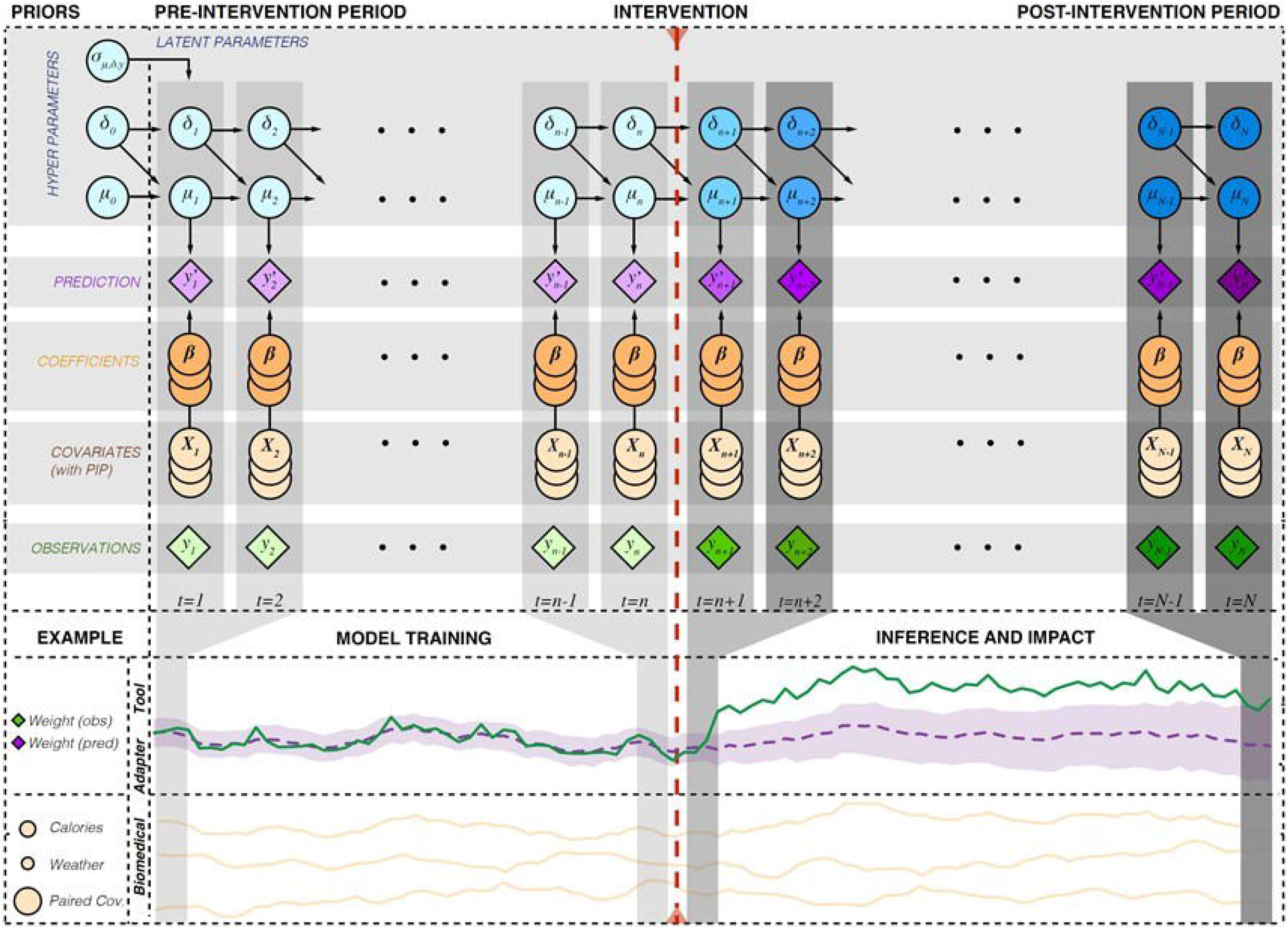
Schematic and illustration ofthe Bayesian structural time series. Latent parameters and hyperparameters are shown in blue, observations are shown in green, covariates in yellow (with a property ofposterior inclusion probability), coefficients in orange, and predictions in purple. Predictions have an associated prediction interval shown in light purple. An illustrative example shows weight over time, with various covariates, being modeled. Covariate posterior inclusion probability is given by the size of the circle. Assuming an intervention ofincreased diet, the model detects a strongimpact on the post-intervention weight.

## STATISTICAL FORMALISMS

### Bayesian Structural Time Series

In order to understand the causal impact of interventions on longitudinal datasets that may be affected by a wide range of factors it is important to have a statistical framework that considers carefully the prior and posterior measurements as well as covariates that may affect the response variable. We detail aspects of this framework below.

We assume for a given time point, *t*, there exists an observation *y_t_*, linked to a variety of other parameters, 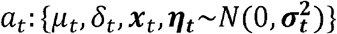. As time progresses from *t* to *t*+1, the other parameters also progress due to their time dependency.

There also exists a set of hyper-parameters 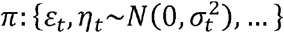 that define the initial conditions of the parameters above. The hyper-parameter *ε_t_* is a prior that reflects previous knowledge about the system and is used for the initial set of parameters in *a_t_*. More specifically, we use the following equations to represent our time series data.

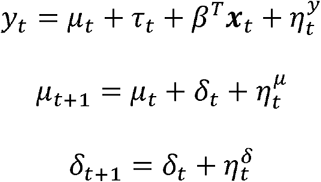

Here we see that the equation involving the observation or response variable, *y_t_*, depends on *μ_t_*, and additionally a variable ***x**_t_* to represent the covariates. Furthermore, we additionally include another layer of dependency, *δ_t_*, which *μ_t_* depends on. Because our observation, *y_t_*, can be biased, it is important to take into account as many covariates in the form of ***x**_t_*. Each equation has an error term represented by *η_t_*. Since each of the processes are different, the error term is also different. For example, the error associated with *μ_t_* is 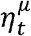. These *η_t_* are drawn from some distribution satisfying 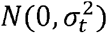. Similar to other regression-based models, the covariates here are scaled by a coefficient vector *β^T^*. The model also employs a spike and slab method to penalize and select for important covariates when the covariate matrix is large. Additionally, one important feature of each covariate is the posterior inclusion probability (PIP), which can be calculated from the sum of all posterior probabilities of all regressions that include that particular covariate (20). The PIP gives a ranking measure to show how favorable the inclusion of a particular covariate is. The variable *μ_t_*, the underlying “state” of the response, contributes to our response variable, and its representation is given as a time dependent equation. If we ignore the *δ_t_* variable, our equation becomes

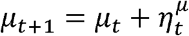

and this is a representation for the random walk. That is to say, in this simplified form of *μ_t_*, our observation is a random walk, where at each time point, there is some progression to the next time point in random fashion, based on the parameter *η_t,μ_*. However, one optimization that can be done for this parameter is the inclusion of *δ_t_*, which results in the pair of equations

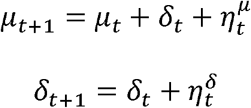

Here *δ_t_* follows a random walk, and the *μ_t_* equation is dependent on *δ_t_*. That is to say, *δ_t_* serves as a trajectory or slope parameter that helps to guide the behavior of *μ_t_*. We can still imagine *μ_t_* as a random walk, but now with some additional trend vector. If *μ_t_* is steadily increasing, it is likely that at *μ*_*t*+1_ there is also an increase, due to the inclusion of *δ_t_*. In other words, the *δ_t_* allows for more stability between each time point and serves a purpose similar to a slope parameter. We provide more examples in Supplemental Figures S1-5.

### Evaluating Intervention Impact

While the above model describes longitudinal sensor data well, it is important to consider how the pre- and post- intervention periods differ. To determine if an intervention was effective in bringing about a change in our dependent measurement, *y_t_*, it is important to have a framework in which comparisons can be quantified. To do so, we define the pre-intervention period as time points *t* = 1, …, *n*, while the post-intervention period is defined as *t* = *n* + 1, …, *N*. Furthermore, we define observations as *y* and predictions of the model as *y*’. Therefore, the set of observations in the pre-intervention period, *y*_1,…,*n*_, serve as the training data, based on all covariates. At each time point *t* = 1, …, *n*, we update our parameter set of the model to better fit *y*_1,…,*n*_ using a Markov Chain Monte Carlo (MCMC) method of sampling from the posterior distribution. After estimation of parameters in the pre-intervention period, we predict the counterfactual in the post-intervention, *y*’_*n*+1,…,*N*_. This is done by using the parameter set defined in the pre-intervention period and the covariates from the post-intervention period. Because *y*’_*n*+1,…,*N*_ has an associated prediction interval, we can calculate a significance p-value associated with the difference between *y*’_*n*+1,…,*N*_ and *y*_*n*+1,…,*N*_. While we can get a p-value at every time point in the post intervention, it is more useful to consider the impact that an intervention had on the whole post-intervention period. The p-value associated with the full post-intervention time period (*t* = *n* + 1, …, *N*) is known as the *cumulative impact*. It should be noted that the prediction interval associated with *y*’_*n*+1,…,*N*_ generally will increase as time progresses. This is one advantage of the Bayesian model and allows for an evolving cumulative impact. This is due to the fact that at every time point after the intervention, the variance associated with the distribution that the prediction is drawn from is compounded at every time point. It is useful to factor in this temporal aspect since the confidence that an intervention resulted in a causal impact may be dependent on time.

### Biomedical Adapter Tool and Software Implementation

We provide a customized biomedical adaptor tool around a specific Google implementation of the Bayesian structural time series framework that uniformly processes, prepares, and registers diverse biomedical data. Specifically, time series data from biosensor data that are measured at different time points and intervals are unified so that given a time interval set *t* = {1, …, *N*}, there exists a set of variables (observation and covariates), *y_t_* and ***x**_t_* for each time point in *t*. The adaptor tool that applies Causal Impact and bsts to evaluate a user defined intervention and gives a report of results (8,21).

## RESULTS

To showcase the wide applicability of the Bayesian structural time series framework for biomedical applications, we provide examples of analysis from various types of sensor data. First, we apply the Bayesian structural time series framework on a toy example where we collected data from an Android phone sensor in order to give intuition for and demonstrate several key features of our model. Second, we apply the model to a real-world example – environmental sensor data collected from an air quality device – and show the usefulness of our model in identifying the effect of simple interventions in real world data. Third, we provide a core biomedical example that showcases a strong application of the framework and the potential it has in personalized medicine. In particular, we collected extensive biosensor data from a diabetes and exercise study which aims to understand how structured exercise can help to stabilize glucose levels throughout the day. The framework was then used to assess the impact of an exercise regimen on clinical health markers of glucose level. We find a strong stabilizing effect on glucose after this exercise intervention. While these data are informative, they currently lack several attributes that we expect future biomedical data to have. In particular, we expect future biomedical datasets to have a large number of context-rich covariates (e.g. activity in different locations or environments) that can dramatically increase intervention assessment. Thus, we give a final example dataset rich in covariates to showcase how these (paired) covariates can be used creatively to more accurately assess the effect of interventions. Specifically, we collected human behavioral sensor data to showcase the patterns of crime across different neighborhoods in a city, and how the introduction of a mobile application aimed at increasing social cohesion affected these patterns. Below, we give more details regarding each of the specific sub-studies and their corresponding findings.

### Toy Example: Google Android sensor

As a toy example to illustrate how the Bayesian structural time series framework works, we used Google Science Journal app on Android and collected various mobile sensor data for a simple experiment involving a person spinning in different scenarios. The goal of this analysis was to demonstrate the flexibility of our model in determining significant causal impact of human physical actions before and after an intervention as well as to provide intuition for the utility of paired covariates at reducing the effects of noise. A schematic of the results is given in Figure 3A, B, and C. First, a single user’s longitudinal data was used to estimate the effect of an intervention (Figure 3A). An example of this could be sleep data and the effect of changing one’s pillow, and where no other data are available to the user. Here we show an individual spinning with a sensor measuring acceleration when held at arm’s length; the intervention is to bring the sensor closer to the body while spinning at the same rate, thereby increasing the acceleration. The model successfully detected the state change associated with the intervention, as shown by the cumulative change relative to the predicted values.

**Figure 3.**
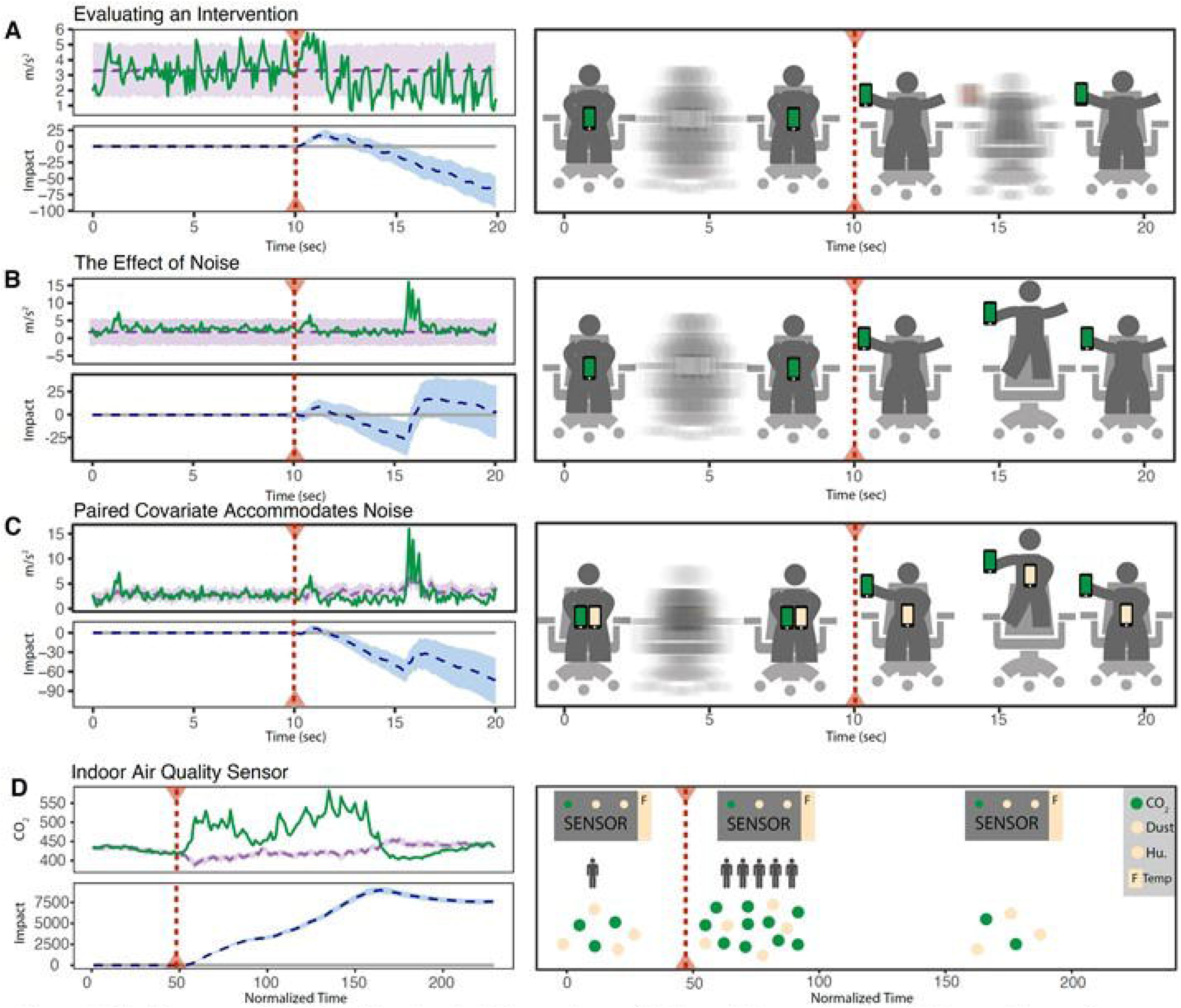
Performance ofthe Bayesian strucrural time series model in model experiments with known interventions. Accelerometer measurements of(A) a person spinning in a chair holding sensor near body and then extended to arm's length at the intervention; **(B)** with simulated “noise” produced by a hop during the intervention period and **(C)** with a paired covariate (second sensor) that is not affected by the intervention but experiences the “noise” hop. D) Using an indoor air quality sensor, CO2 is measured with a variety of other covariates. The intervention (arrival of office members) causes an increase in CO2 and is detennined to be impactful using the model.

Next, we demonstrate how the previous scenario is susceptible to noise, i.e. something that affects the signal but is not related to the intervention. In this case the noise is represented by a hop, which occurred after the intervention began and was therefore not predicted by the model (Figure 3B). In this case the model does not detect an effect of the intervention – the hop affected the signal to such a degree that the confidence in the prediction decreased, i.e. the credible intervals widened greatly. To continue the sleep study analogy mentioned above, the noise could be loud neighbors moving in next door (adding noise in a literal sense), which leads to poor sleep in a way that is unrelated to the pillow intervention.

Finally, we show how including a paired covariate -- another data stream that affects the state being measured but is not related to the intervention – can effectively correct for noise (Figure 3C). In this case two sensors are held while spinning, but only one is subject to the intervention (brought close to the chest). Both, however, experience the noise (hop), which the model is then able to control and correctly identifies the intervention like the case without noise (Figure 3A). In the sleep analogy, an effective paired covariate could be another device measuring the sleep of one’s partner sharing the same bed, but who did not change their pillow. This is distinct from a non-paired covariate which could be similarly effective, e.g. a device measuring the decibel level in the room; it is relevant to the state (sleep quality), is not affected by the intervention (pillow change) and identifies the noise that affects the state (loud neighbors). However, a paired covariate has the characteristic that it is able to control for unknown confounders, akin to a control arm of a randomized controlled trial.

### Simple Real-world Example: Environmental Sensor Data

We then transitioned to a simple real-world example where we performed a similar analysis using data collected from an AWAIR monitor, with measurements of CO_2_, dust, humidity, and temperature (Figure 1E). The data were collected over a one-month period, with measurements taken every 15 minutes. The data were aggregated at each time point across all 30 days to give a smoother signal and limit the analysis to one intervention with a clearly delineated a pre- and post- intervention period, namely exposure of the room to people. It should be noted that though intervention often implies some treatment put in place, it can be widely adapted in the realm of Bayesian structural time series to allow for any disruption or change in status such as the effect that people have on environmental CO_2_ levels. After correcting for covariates such as dust, humidity, and temperature (22), there was a significant increase in CO_2_ in the hours when workers were in in the room with the sensor (p-value < 0.001). Figure 3D shows the CO_2_ measurements across the aggregated time points in a 24-hour time frame. We can see that as CO_2_ levels increased drastically after the 9am time point, the cumulative causal impact in the post-intervention period increased. This cumulative impact increases until around 5pm in the evening, where the cumulative impact tapered off, signaling a state was reached in the post-intervention that was similar to that of the pre-intervention period. Although there is one p-value associated with the whole post-intervention period, this framework can show confidence intervals for each time point in the post-intervention period as a function of both the observation and covariates. This helps us to define period of time where the intervention is most effective.

### Core Biomedical Example: Clinical Sensor Data

We next provide a core biomedical example of how the framework can be applied to clinical sensor data. In particular, we collected biosensor data from a person with type 1 diabetes over a 12-week period who completed an exercise regimen (intervention). We chose to model this set of biomedical data because it is arguably one of the most established applications of personalized medicine today (23,24). People with type 1 diabetes are advised to intensively manage their blood glucose to maintain it within the target range. Since this process is quite involved and requires continuous adjustments based on numerous biological factors, the result is a complex and high-stakes problem in personalized medicine.

Figure 4A shows the data from a continuous glucose sensor and insulin pump as well as a comprehensive set of Apple Watch data from the study. The 12-week evaluation spanned an initial 2-week sedentary period followed by a 10-week exercise regimen. The Apple Watch and the insulin pump used by the patient provided several potential covariates, which could help us understand the glucose sensor data. We took glucose readings from the participant’s glucose sensor and aggregated them into 24-hour values by transforming the values into two clinically relevant indicators of glucose stability, percent-in-target and percent-above-target (25). In general, values above the target range are predictive of long-term organ damage, while values below the target range lead to acute hypoglycemia (26), a state of low blood glucose with symptoms that can range from mild fatigue and confusion to life-threatening coma, posing immediate threats to safety as well as long-term psychological consequences (e.g., fear of hypoglycemia (27)).

**Figure 4.**
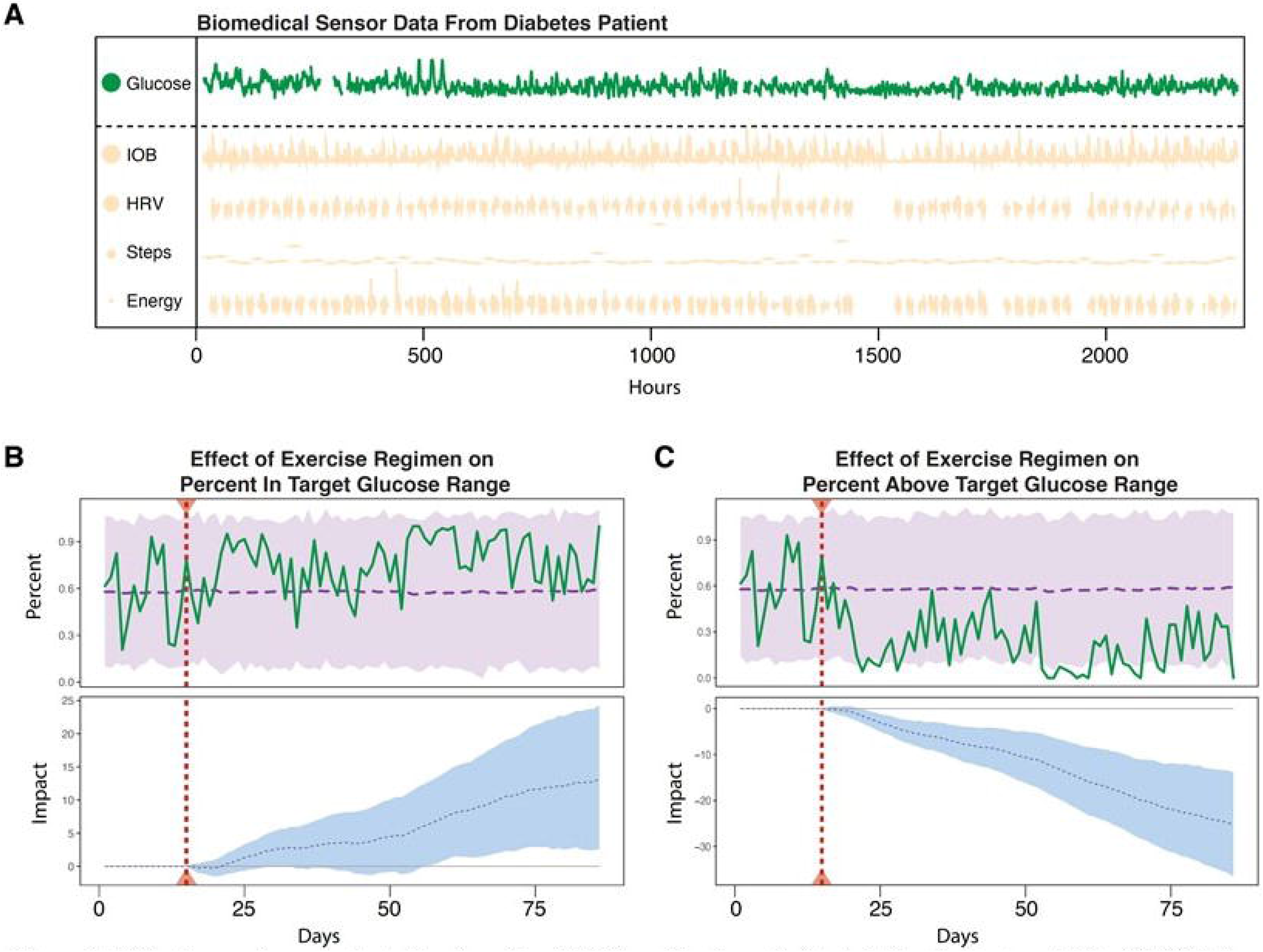
A) Continuous glucose, estimated insulin on board (IOB), and Apple watch data,including heart rate variability (HRV), daily steps taken, and energy expended, over 12 weeks for the participant ofinterest. B) Analysis of a 10-week exercise regimen's causal impact on the percentage ofdaily glucose readings in the target range (percent-in-target). C) Analysis of a 10-week exercise regimen's causal impact on the percentage ofdaily glucose readings above the target range (percent-above-target). HRV,heart rate variability. IOB, insulin on board.

Maintenance of glucose levels in the target range is achieved by strategically managing a triad of factors: insulin administration, diet, and exercise. The timing and dosing of insulin (which decreases blood glucose) with carbohydrate ingestion (which increases blood glucose) must be carefully balanced. The role of exercise, however, is less clearly defined because it can either decrease or increase blood glucose during and up to 24hr after a session. Determinants of the direction and magnitude of the glycemic response to exercise are numerous and include **1)** exercise characteristics (high vs. moderate or low intensity), **2)** individual characteristics (endogenous insulin sensitivity, and the effect of physical fitness to increase insulin sensitivity), and **3)** contextual factors (pre-exercise blood glucose level, insulin- and carbohydrates-on-board, and concentration of counter regulatory hormones) (23).

Using our Bayesian structural time series framework, we find that the exercise regimen was effective in increasing the percent-in-target and decreasing the percent-above-target (*p* = 0.014 and 0.002 respectively). These results are shown in Figure 4B and 4C. Thus, our model supports the existence of a positive causal effect of this particular exercise regimen on maintaining a healthy glucose level for this individual. Furthermore, of all the covariates we used, insulin on board (IOB) was found to have the highest posterior inclusion probability, which is a ranking measure used to see how favorable the inclusion of a variable is in the model. This is reasonable since insulin is essential to glucose control.

### Example with Data Rich in Covariates: Human Behavioral Data

In our final example, we demonstrate how data rich in covariates can significantly improve assessments of intervention impact. While the example we provide is not clinical in the traditional sense, we provide this example as a way to showcase how the Bayesian structural time series framework performs when given extensive data and paired covariates – an important consideration for future biomedical data. One aspect of a rich covariate set could be location information, an important factor in many clinical or personal health applications. For example, many watches collect GPS tracks of a run as well as steps and heart rate (e.g., Apple Watch, Garmin Forerunner). The Bayesian structural time series framework can flexibly accommodate location data through the inclusion of spatial correlation matrices. In addition, the spatial data can be used to segment the data where the intervention does and does not have an effect. By this method one can create “synthetic” paired-covariates, in that data are from different spatial segments from the same source. However, spatial data are often protected for privacy concerns and are therefore less accessible to researchers. We therefore demonstrate the utility of the framework on more accessible behavioral data that share many characteristics: the effect of a social monitoring application called SeeClickFix on negative behavioral patterns (crime).

Crime patterns show similar characteristics to spatial mobile health data. They are affected by covariates such as temperature and precipitation (28) and can be linked to many other features such as census data of household incomes or education (29) (analogous to covariates of movement such as age, physical fitness, etc. (30)). SeeClickFix is a smartphone and web application developed to allow users to report issues in their communities including non-violent crimes. Posts can be voted on and supported by other users’ comments and local government agencies acknowledge issues and post when they have been addressed. It has been hypothesized that SeeClickFix and similar tools may reduce crime through establishing social cohesion, promoting collective efficacy (31).

Following the intervention (creation of SeeClickFix) there was no detectable decrease in crime across the entire area (Figure 5A), nor in particular neighborhoods (Figure 5B). However, one would expect many other factors besides the introduction of SeeClickFix to affect crime in this time (e.g. increases in the police force, changes to local employment, city-wide initiatives). The Bayesian structural time series framework can leverage the spatial information in these behavioral data to search for a paired-covariate, in this case referring to locations not affected by the intervention (no SeeClickFix use) in order to control for other, unobserved effects on the outcome variable (increases in the city police force). We aggregated crimes and SeeClickFix posts by neighborhood (Figure 5C) and then modeled used a neighborhood that was not affected by the intervention as a control (i.e. a neighborhood without SeeClickFix posts), but that would experience any city-wide initiatives that might affect crime (Figure 5D). Neighborhoods with heavy SeeClickFix use showed an effect of the intervention on crime when controlling for unobserved factors with synthetic paired covariates (Figure 5E).

**Figure 5.**
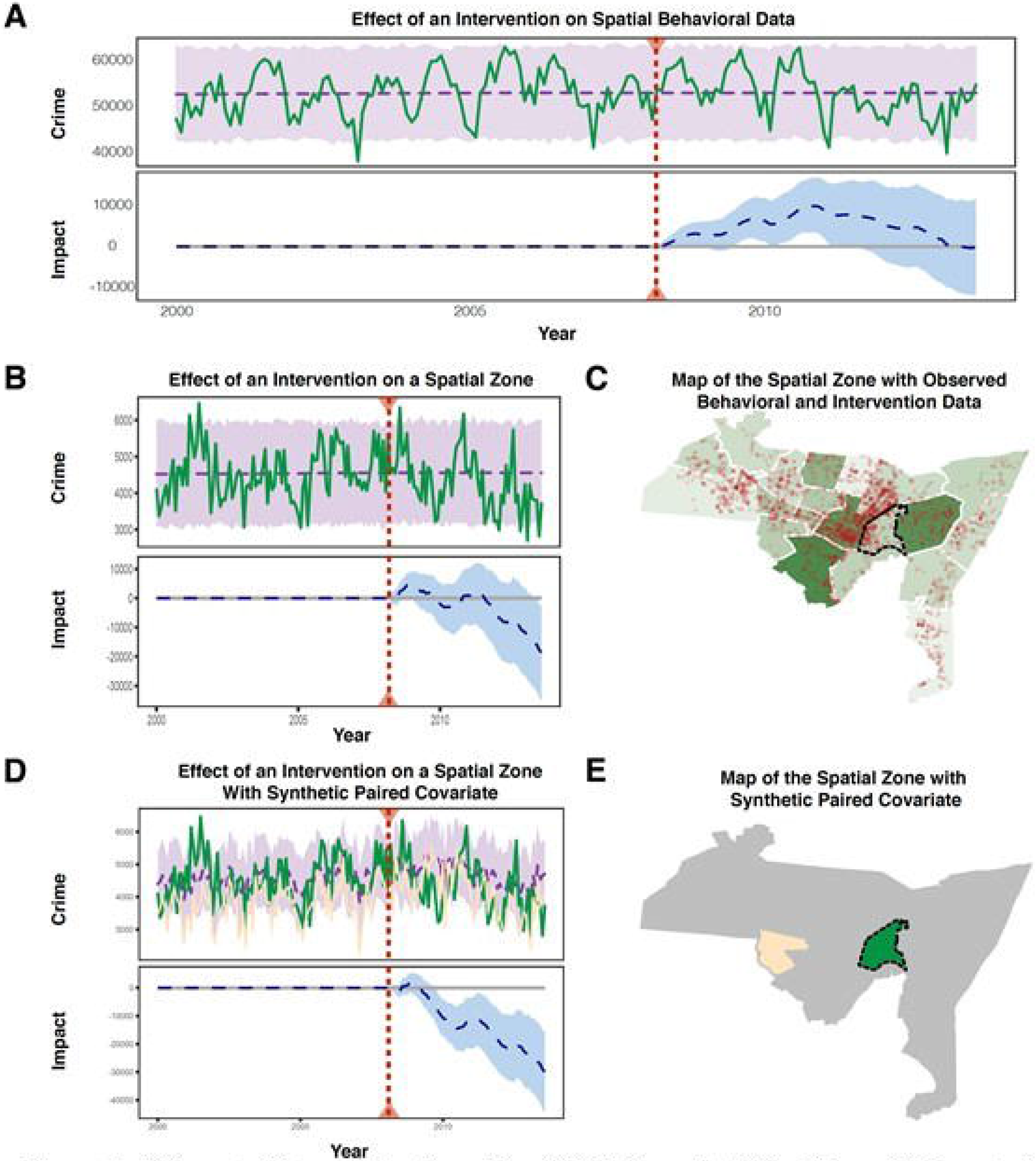
A) Impact ofintervention (use ofSeeClickFix) on all ofNew Haven. B) Impact of intervention on only one neighborhood, Wooster Square. C) Spatial data ofNew Haven showing variety ofSeeClickFix usage throughout various neighborhoods. D) Impact of intervention on Wooster Square crime using West River as a paired covariate. E) Illustration showing the observed data, Wooster Square, versus the paired covariate, West River.

## DISCUSSION

In this paper, we demonstrated how the Bayesian structural time series framework can be applied to biomedical sensor data and personalized medicine as well as created a wrapper software to facilitate the framework’s use. We additionally showcase how a rich dataset with complex covariates can benefit from the framework. While some studies demonstrating the success that linear models have on understanding such data do exist (18,19,32–34), there is a lack of emphasis on considerations for the temporal aspect of interventions and how effective interventions are at different time points in the post-intervention period. Linear modeling frameworks lack the flexibility to evaluate the intervention’s effect strength at all points in the post-intervention period, while existing Bayesian frameworks forecast based upon predictors (e.g., forecast continuous blood glucose based upon covariates like the ones we obtained from insulin and smartwatch devices) but do not evaluate deviation from this forecast following an intervention of known timing and duration to determine its impact (35–37). In this study, we illustrate the benefit of using a Bayesian structural time series model in modeling the behavior of various longitudinal data collected via apps and sensors. These data demonstrate properties that are commonly found across other wearable sensor data, which are increasingly gaining in popularity (3). In order to effectively integrate such data with personalized medicine and health in the future, we must understand how specific timings and types of interventions impact individual patients (2–4).

It is also important to consider the contexts in which this modeling framework does not yield significant advantage over a linear framework. A stable and consistent intervention effect over the entire post-intervention period should be evaluated equally well by linear models. Given the computational complexity of the Bayesian method due to MCMC sampling for each time point, such considerations could be very important for large datasets. Furthermore, due to the nature of this type of model and the slight degree of randomness in parameter estimation, it is possible to have varying results in calculating the impact of the intervention. This is in contrast to the results found from linear models, which generally demonstrate a singular solution.

We show that the Bayesian modeling framework can take into account the rich covariates in our behavioral sensor dataset – specifically the paired covariate structure between different locations – and give results that are unbiased. Furthermore, the framework also considers the temporal properties, ensuring that predictions of intervention impact are conditional on duration of the intervention. This is especially important for future data from sensors and mobile health sources as one of the major concerns of personalized medicine is ensuring the right time and duration of a treatment or intervention is considered (5). With time varying confidence intervals, we can begin to understand not only the effectiveness of an intervention but determine the most effective intervention plan necessary to reach a desired result.

When applying this method to various datasets, an additional consideration is how informative the covariates are. For example, in the case of the sensor data derived from the diabetes study, even though we were able to isolate the effect of the 10-week exercise regimen upon both metrics of glucose stability, more covariates could be used in the future to improve the prediction. Similar biological studies in the future could benefit from obtaining other clinically relevant covariates such as plasma cortisol, epinephrine, growth hormone, glucagon, and directly sensed (not estimated) insulin levels or more to use high precision sensors with reduced noise. Furthermore, paired covariates (e.g. control group) should be taken into consideration when designing studies, as they can greatly increase model accuracy. While it is true that even an improved set of covariates could lead to more accurate results, we showcase here that the framework we have now is able to demonstrate results that converge with current literature showing favorable impact of exercise upon blood glucose control metrics (38). We also note that one assumption of our model is that the intervention does not affect any of the covariates in the post-intervention period. For example, though the exercise regimen is aimed at stabilizing glucose levels, we note that exercise and physical activity can have a general effect on various biological factors within the human body, including some of the covariates used. We minimize the effect of this through our transformation of variables and aggregation on a 24-hour block.

The Bayesian structural time series framework is a flexible modeling approach that, with relatively minor specialization we provide through our tool, can be applied to diverse biosensor data. It has features that few methods in biomedicine share, such as dynamic forecasts to observe the effect of an intervention and its evolution over time. These features are critical to advancing personalized medicine and realizing the challenge of relating genotype and phenotype data in the context of human research.

## METHODS AND MATERIALS

### Data Collection

There exist many datasets similar to those of mobile sensors and we should not confine ourselves to just the traditional accelerometer and gyroscope data that one would traditionally think of when envisioning sensor data. In studying the causal impact of an intervention, we show that most data sharing longitudinal qualities allow for development of algorithms and exploring how such algorithms can be useful in analyzing various sensor data. The data we analyze in this paper consists of environmental sensor data, physical activity sensor data and human behavioral sensor data.

Due to the longitudinal nature of sensor and wearable data, these data not only demonstrate interesting patterns in a response variable, but also are closely tied to the temporal property of a phenomena. By introducing the aspect of time, it becomes important to find models that leverage temporal considerations and use them to make accurate assessments about the data. Also, some of the data showcase complex covariates (paired) and can be used to better correct for unknown and hidden biases. More details about the data collection and data types are given below.

#### Google science journal data collection

Data were collected using the linear accelerometer measurement function in the Google Science Journal application on a Samsung Galaxy S8 smartphone. Each experiment lasted 20 seconds. One or two instruments were held at arm’s length while spinning at a constant rate for 10 seconds. Next, one instrument was brought in close to the body for 10 seconds while maintaining the spinning rate. In the case of the noise simulation, the experimenter hopped once after 15 seconds (halfway through the intervention period). Data were exported from Science Journal and analyzed using the CausalImpact package in R.

#### AWAIR Collection

Data were collected from a one-month period from the AWAIR device. The device was placed in an office lab setting where individuals frequented on a daily basis. CO_2_ levels were measured in units of ppm. AWAIR also measures dust, temperature and humidity, which were used as covariates.

#### SeeClickFix Data collection

The application SeeClickFix is a smartphone and web application developed in New Haven, Connecticut, where users report issues in their communities including non-violent crimes. SeeClickFix posts can be supported and commented on by other users, and local government agencies acknowledge and address issues. The SeeClickFix data are publicly available, providing a rich longitudinal and spatial dataset for monitoring behavior and interactions with other users and city representatives. Posts were aggregated by month for the New Haven metropolitan area and by neighborhood (n = 19) from 2007-2015.

Aggregated crime data were shared through a memorandum of understanding with the New Haven Police Department for 2000-2013. Rates were calculated using the 2014 ACS 5-year population estimates (crimes / 10K population per unit area).

#### Diabetes Data collection

We used data from one participant in a single-group clinical trial that was evaluating an exercise intervention for previously sedentary adults with type 1 diabetes. Participants completed a 2-week baseline period then a 10-week exercise intervention, while wearing sensors that continuously monitored blood glucose, heart rate, heart rate variability and physical activity. They continued their normal prescribed insulin therapy, and shared device-recorded dosing logs with the research team. Besides these continuous measures, we assessed chronic diabetes control at the beginning of the baseline period and the end of the 10-week intervention using blood glycosylated hemoglobin concentrations. The 10-week intervention included motivational enhancement of exercise (i.e., patient-centered exercise coaching including instructional videos) and health feedback from biosensors (full details available on clinicaltrials.gov, NCT04204733). Both were delivered through a customized mobile digital application and supported by a coach internally certified in exercise for diabetes (GlucoseZone™, Fitscript^LLC^, New Haven, CT). The study was approved and overseen by the Yale University Institutional Review Board, and all participants provided written informed consent.

##### Biodata Collection

1) 24hr blood glucose was measured every 5 minutes by the Dexcom G6 continuous glucose monitor (San Diego, CA), a subcutaneous wire sensor sampling interstitial fluid glucose content which is converted to estimated blood glucose (validated against venous blood glucose with mean absolute relative difference 9%) (39). 2). Insulin was delivered according to each participant’s usual prescribed therapy. The participant in this manuscript received lispro insulin via the Tandem t-slim Control IQ pump (San Diego, CA). The pump subcutaneously infuses insulin every 5 minutes according to the patient’s individualized settings and current blood glucose levels using proprietary algorithms (40). The patient can also manually dose insulin or adjust some standard settings for meals or other disturbances (e.g., planned exercise). Infusion doses are recorded, uploaded to a central server for exporting and analysis, and converted to estimated insulin on board by the manufacturer’s proprietary pharmacokinetics algorithm. 3) Heart rate (beats per minute, validated against electrocardiography with mean absolute percentage error 1.1%-6.7%) (41) and heart rate variability (standard deviation of interbeat intervals, validated against electrocardiography with intraclass correlation coefficient 0.98) (42) were measured by the Apple Watch 3 (Cupertino, CA) using photoplethysmography. 4) Physical activity was measured by the Apple Watch 3 using accelerometry and converted to kcals per day (validated against calorimetry with mean absolute percentage error ~40%) (43). 5) Diabetes control was measured by glycosylated hemoglobin (DCA Vantage Analyzer (Bayer, Tarrytown, NY) at baseline and PTS Diagnostics A1cNow+ (Indianapolis, IN) at 10 weeks).

##### Clinical Outcomes

The participant was a 63-year-old white non-Hispanic female with type 1 diabetes for 50 years, receiving 81 units/day of insulin (0.9 units/kg body weight/day) and performing no regular exercise at baseline. During the 10-week intervention she received 93 units/day of insulin (1.1 units/kg body weight/day) and exercised on average 2.5 days per week, 26 minutes per session, at easy to moderate intensity (Borg rating of perceived exertion 2.5 / 10). The exercise routines were dynamic, interval-based, and equally emphasized all major muscle groups. Her chronic diabetes control indicated by glycosylated hemoglobin improved from 8.6% at baseline to 6.9% at 10 weeks, achieving the recommended target of less than 7.0% (44). Processing and analysis of all data was done in R and Python and our packaged tool can be found at https://github.com/gersteinlab/scf.

## ACKNOWLEDGEMENTS

This work was funded by the National Institutes of Health (NIH).

GA was supported by a fellowship from the Office of Academic Affiliations at the United States Veterans Health Administration.

