## Supplemental Information for "Bayesian Structural Time Series for Biomedical Sensor Data: A Flexible Modeling Framework for Evaluating Interventions"

Supplemental Figures

Figure S 4 Using the Bayesian structural time series model on the generated data	4

### *Section 1. Additional Statistical Details about the Bayesian Structural Time Series Model*

The Bayesian structural time series model is formulated as below

$$y_{t}=\mu_{t}+\tau_{t}+\beta^{T}\boldsymbol{x}_{t}+\eta_{t}^{y}$$

$$\mu_{t+1}=\mu_{t}+\delta_{t}+\eta_{t}^{\mu}$$

$$\delta_{t+1}=\delta_{t}+\eta_{t}^{\delta}$$

However, it is important to point out that there are additional parameters that could be included but were left out for this analysis. One such important parameter could be

$$\tau_{t+1}=-\sum_{s=1}^{S-1} \tau_{t}+\eta_{t,\tau}$$

This $\tau$ parameter could be considered as a seasonality parameter and can be used to delineate differences observed in a response variable due to seasonal changes. Here $S$ represents the total number of seasons.

### *Section 2. Additional Graphical Examples of the Bayesian structural time series and Causal Impact*

Figure S 1 Generated Data

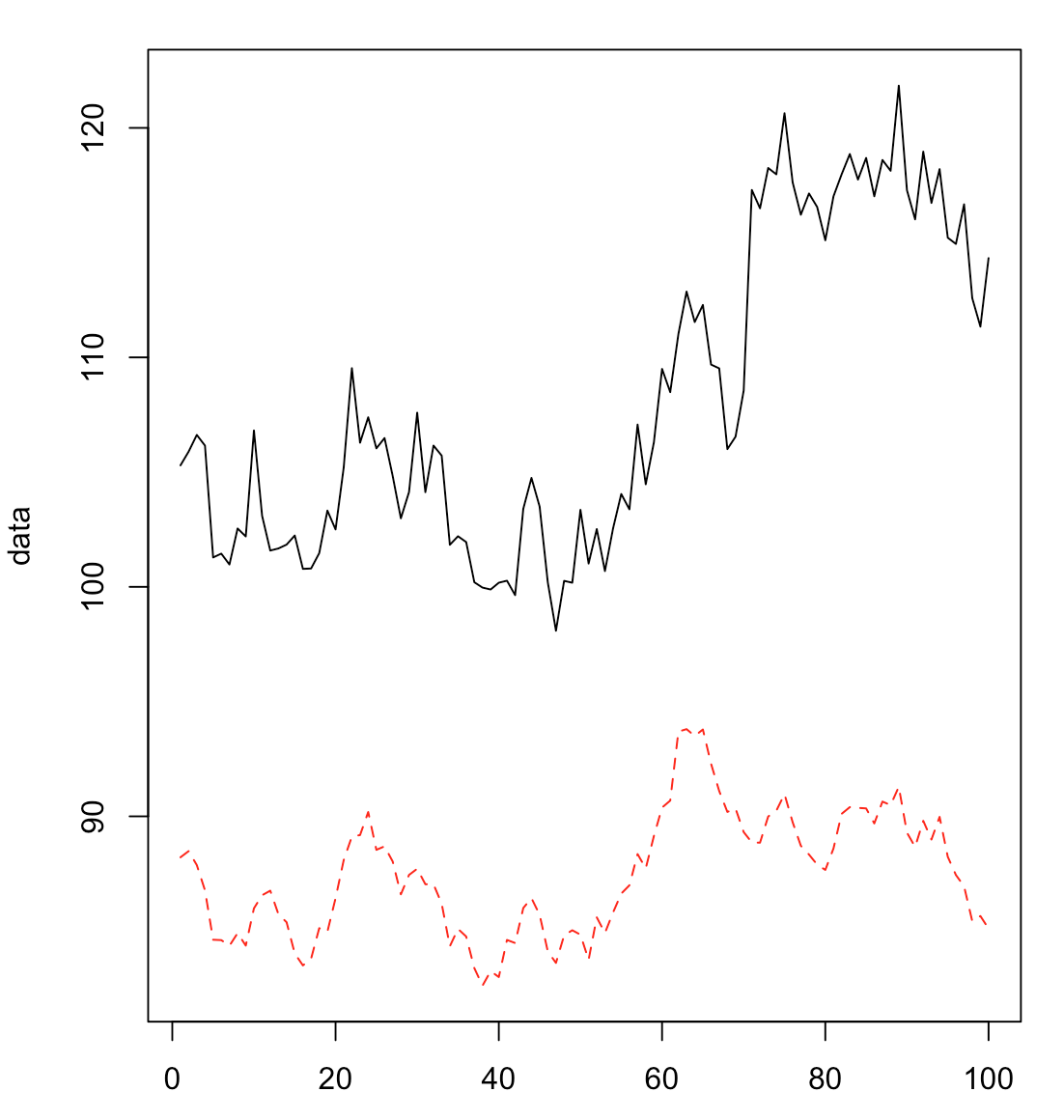

In order to demonstrate the utility of our model on generated data, we use the example data above. The black line shows our response variable and the red dotted line shows a covariate. We assume the intervention occurs at time point 70. In the following panels below, a model is fitted to the pre-intervention period and the counterfactual is predicted in the post-intervention period. The first panel shows the model training and prediction, the second panel shows the pointwise differences, and the third panel shows the cumulative differences. The figures are generated using the CausalImpact package in R by Broderson *et al* 2015.

Figure S 2 Using a linear model with no covariates to model the generated data.

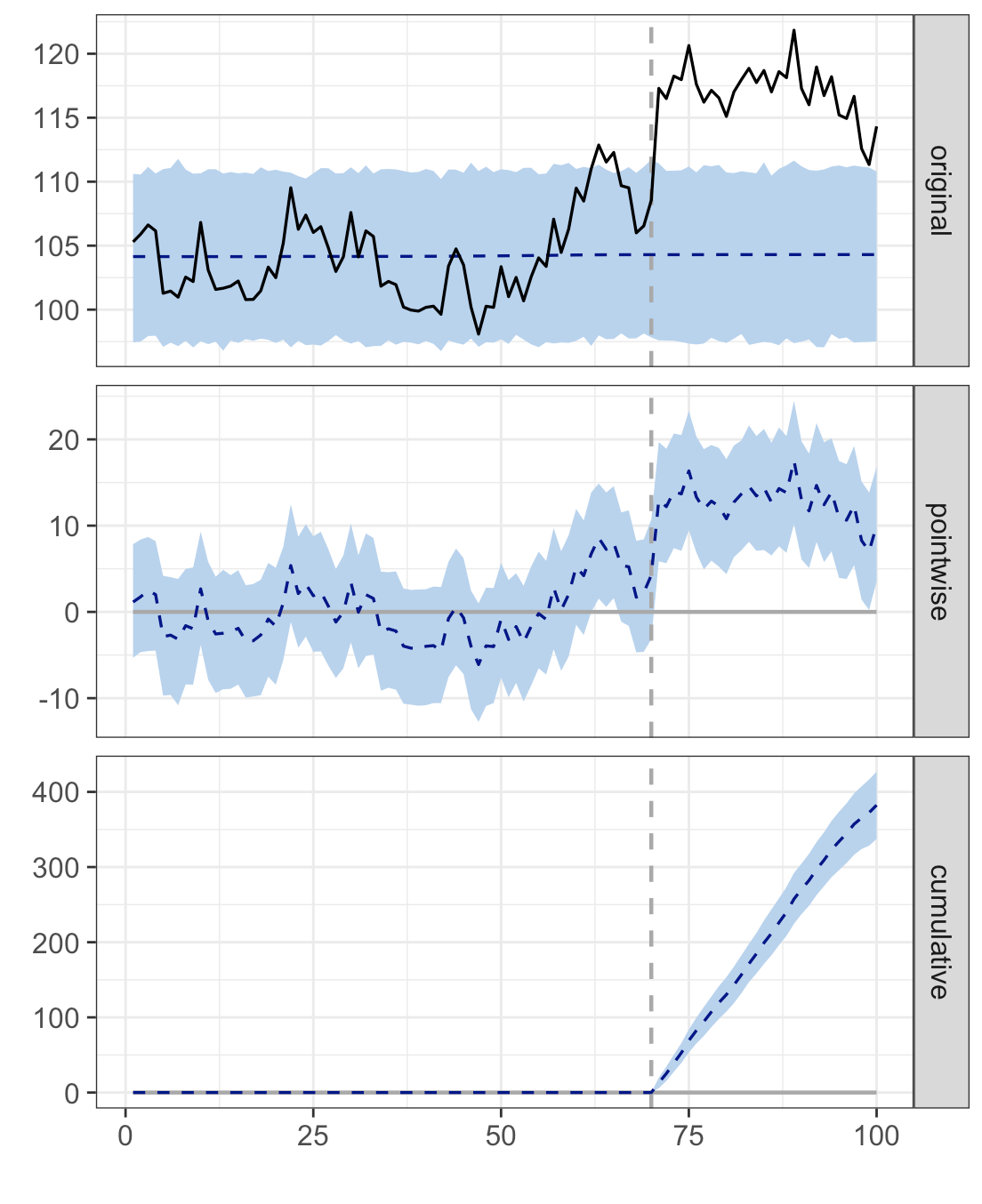

In Figure S2, we apply a linear model to the pre-intervention period, assuming no covariates. As expected, this provides a straight line prediction equivalent to the mean of the pre-intervention data. We can see the line fits the pre-intervention data in the training set very poorly as shown by the confidence interval that extends well beyond the range of the pre-intervention data.

Figure S 3 Using a linear model with covariates to model the generated data.

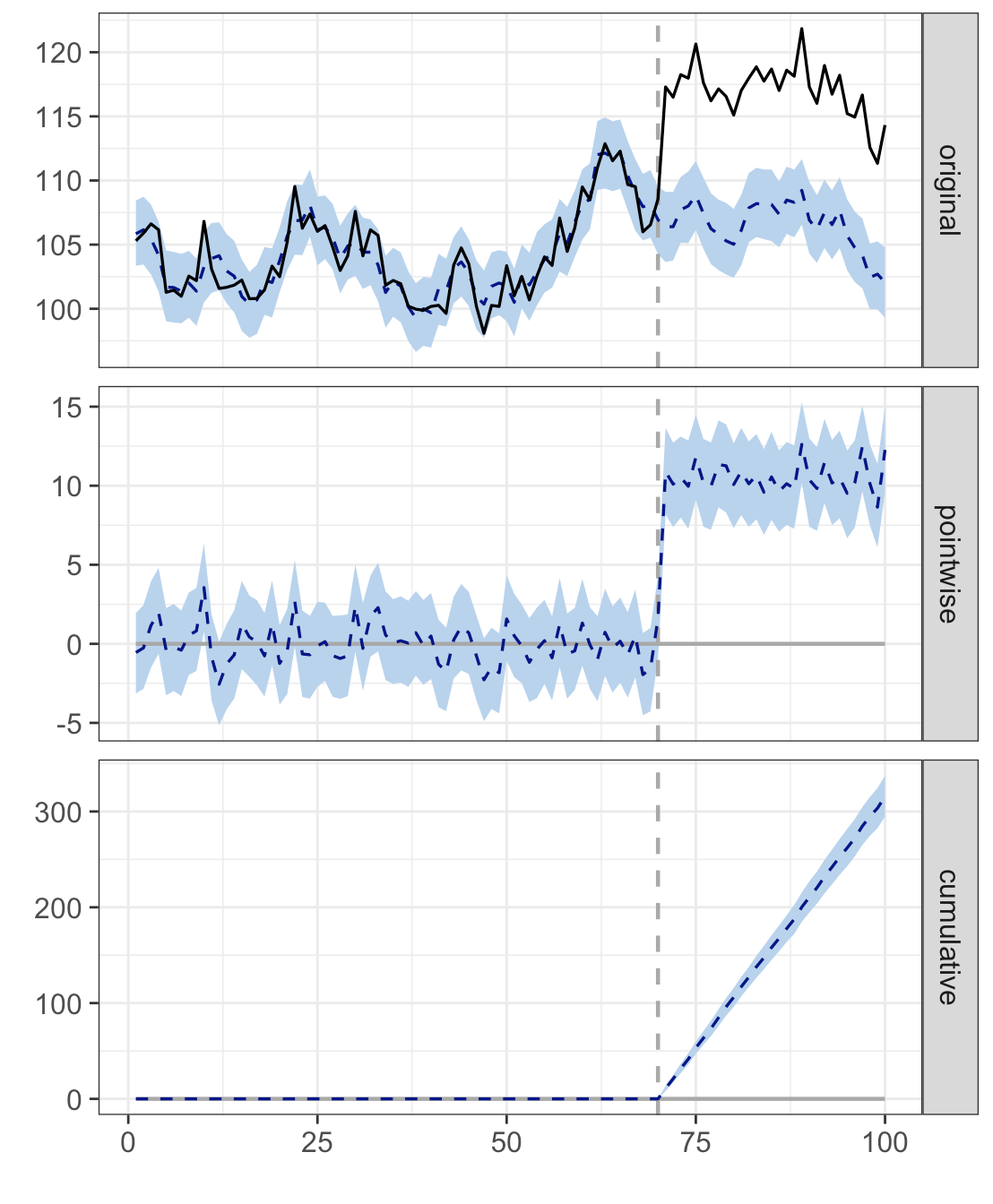

In Figure S3, we demonstrate the use of a linear model with covariates. As shown above, the model is much better at modeling the data in the pre-intervention, but is only able to do so with dependency on covariates, suggesting a high reliance on such data.

Figure S 4 Using the Bayesian structural time series model on the generated data

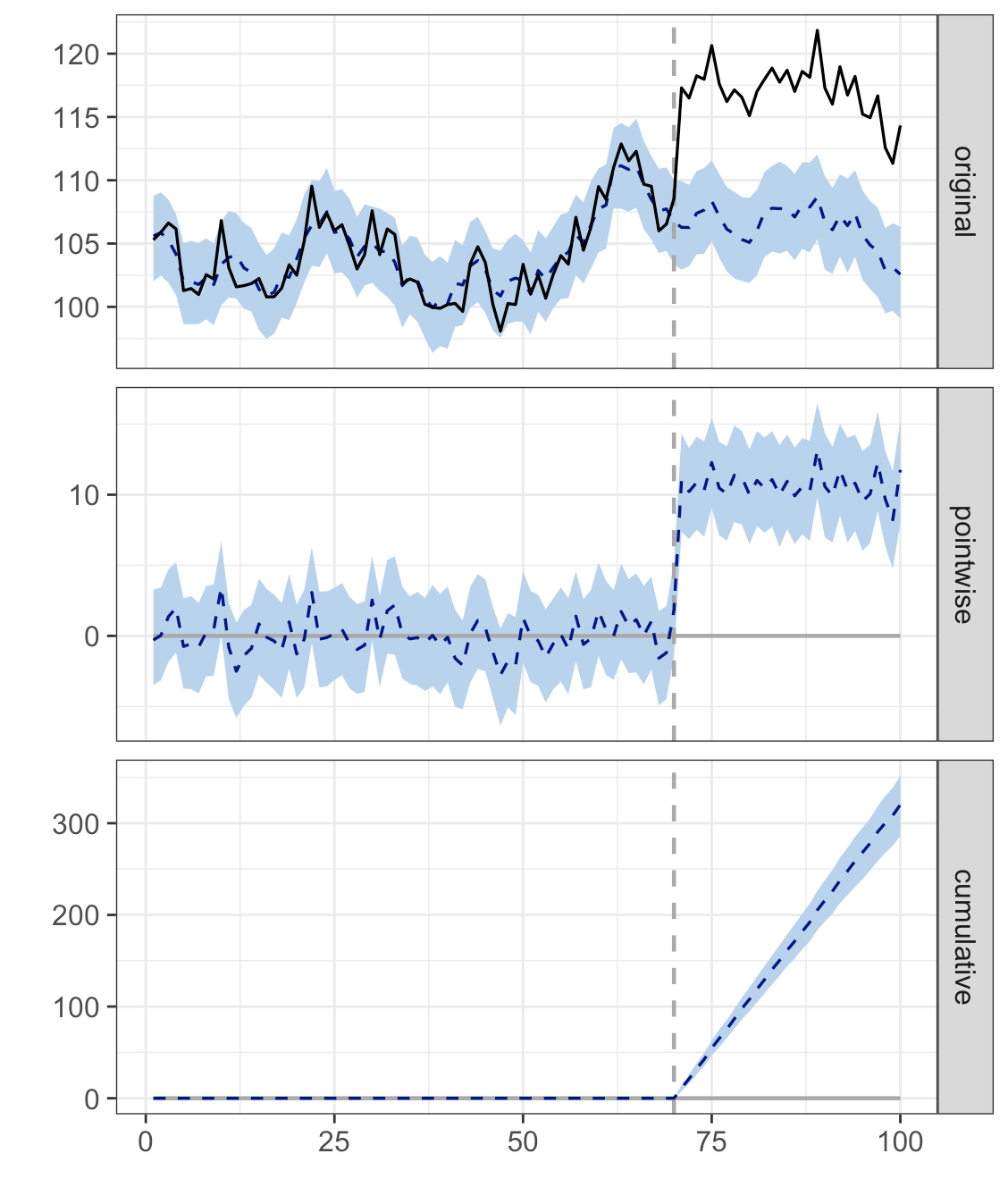

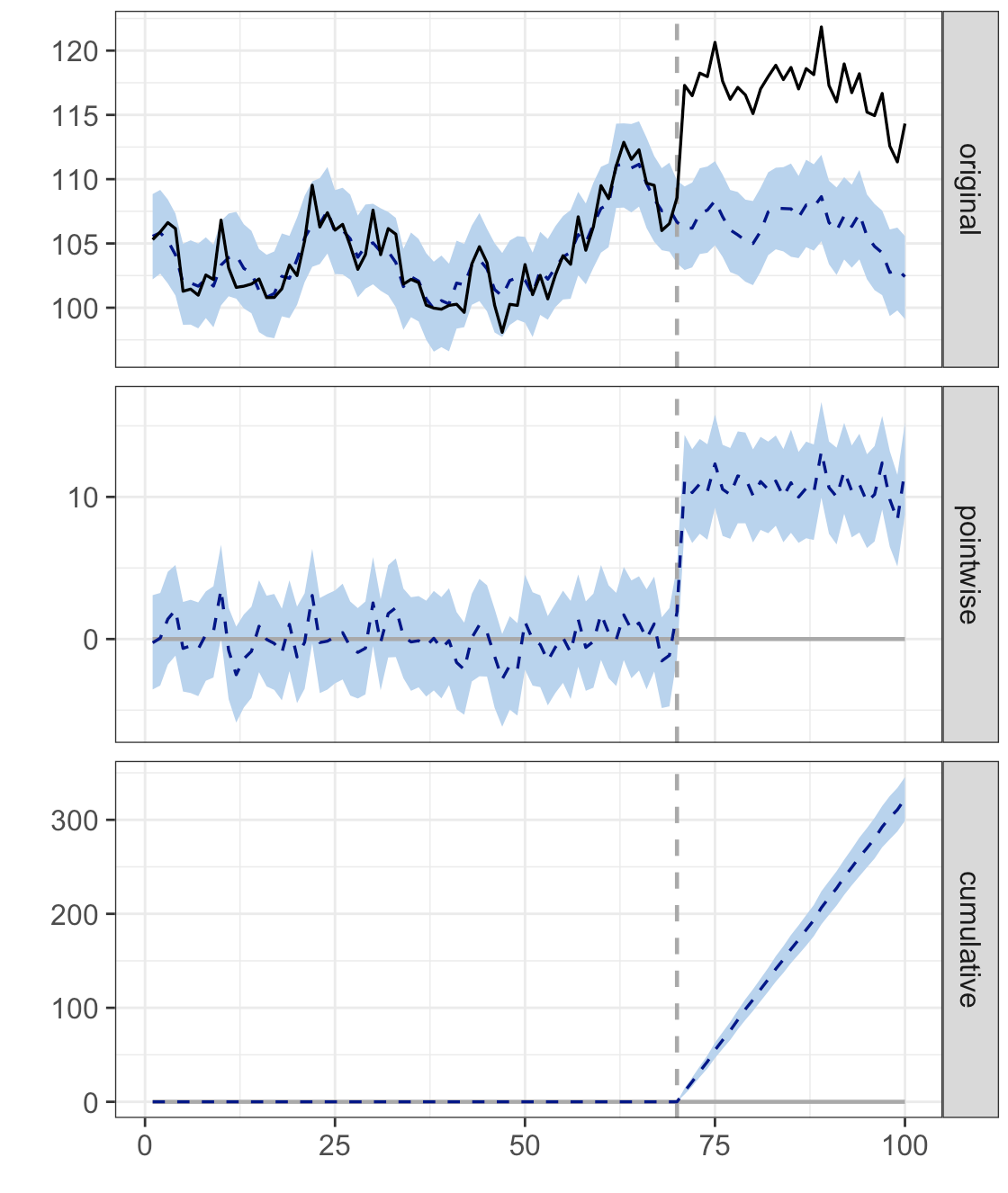

The panel on the left in Figure S4 shows the Bayesian model with inclusion of a $\mu$ parameter while the panel on the right shows the model with inclusion of a $\mu$ and $\delta$ parameter. There is a very slight change in stability of the model as the inclusion of the $\delta$ parameter allows for a slight decrease in the range of the prediction interval, suggesting higher confidence through inclusion of this parameter.

Figure S 5 Using the Bayesian structural time series with no covariates.

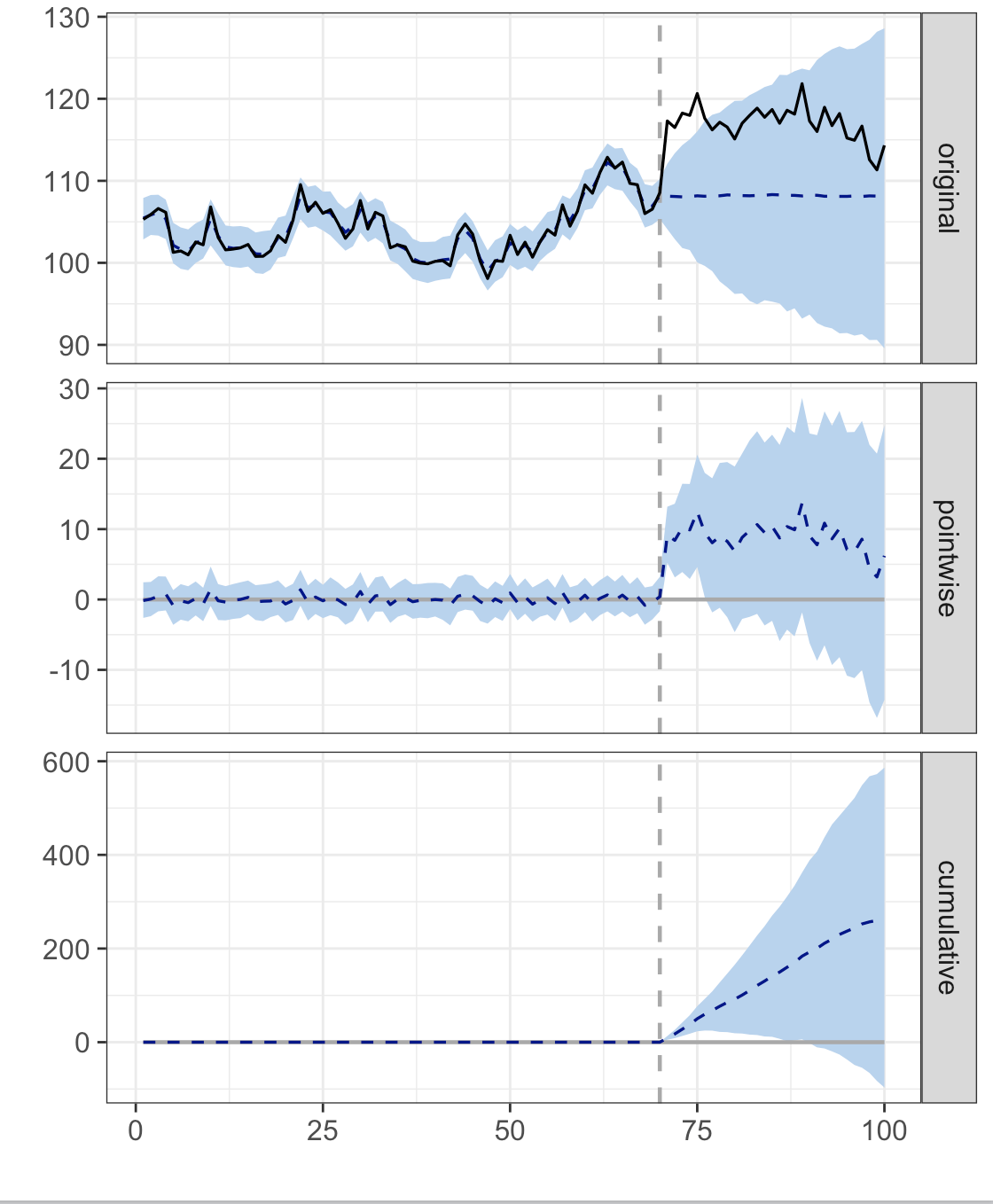

It is often impossible to gather all covariates of a particular response variable. Sometimes it is possible to use paired covariates as described in the manuscript, but other times no covariates exist. Here we show that even without covariates the Bayesian structural time series model is flexible enough to measure the impact of the intervention at time 70, but remains conservative enough by including an evolving and growing prediction interval as to not falsely identify impact long into the future after the intervention.

### *Section 3. Additional Information Regarding the Behavior (Crime Data) Analysis*

Using spatial information is an important part of the analysis of behavioral data in the manuscript. Below we show specific spatial information that can be useful to determine what covariates to use.

Figure S 6 Spatial information from human behavior data set

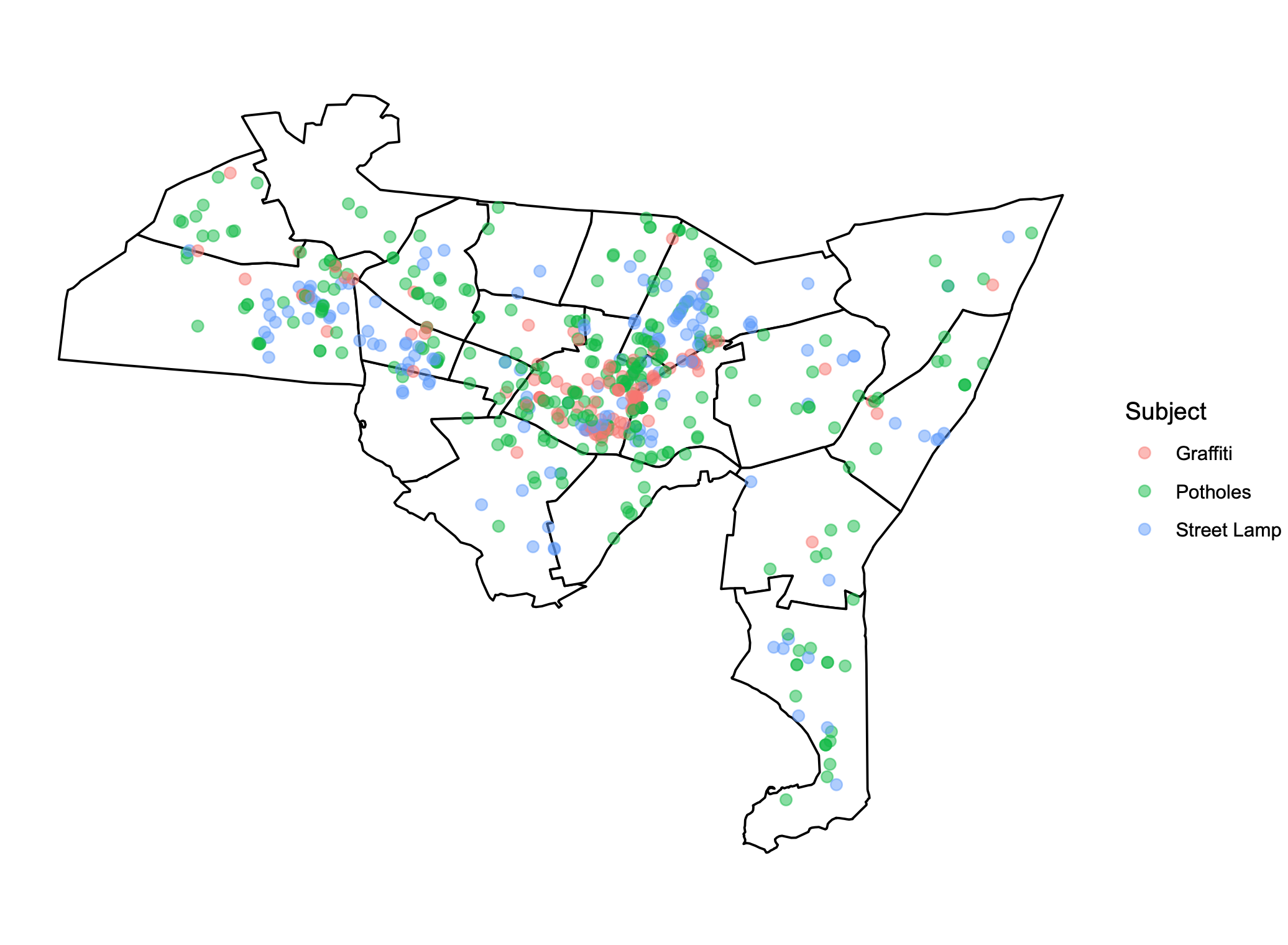

We also include the full output of the analysis of the New Haven crime data (with and without a paired covariate) below. We can see that in Figure S7, the use of no paired covariates makes it very hard to determine how effective the intervention was. We see in the third panel no clear signal of a significant impact. However, in Figure S8, we see that the model clearly is able to determine a signal in the post-intervention, suggesting a clear trend of impact.

Figure S 7 Determining causal impact on behavior data with no paired covariates

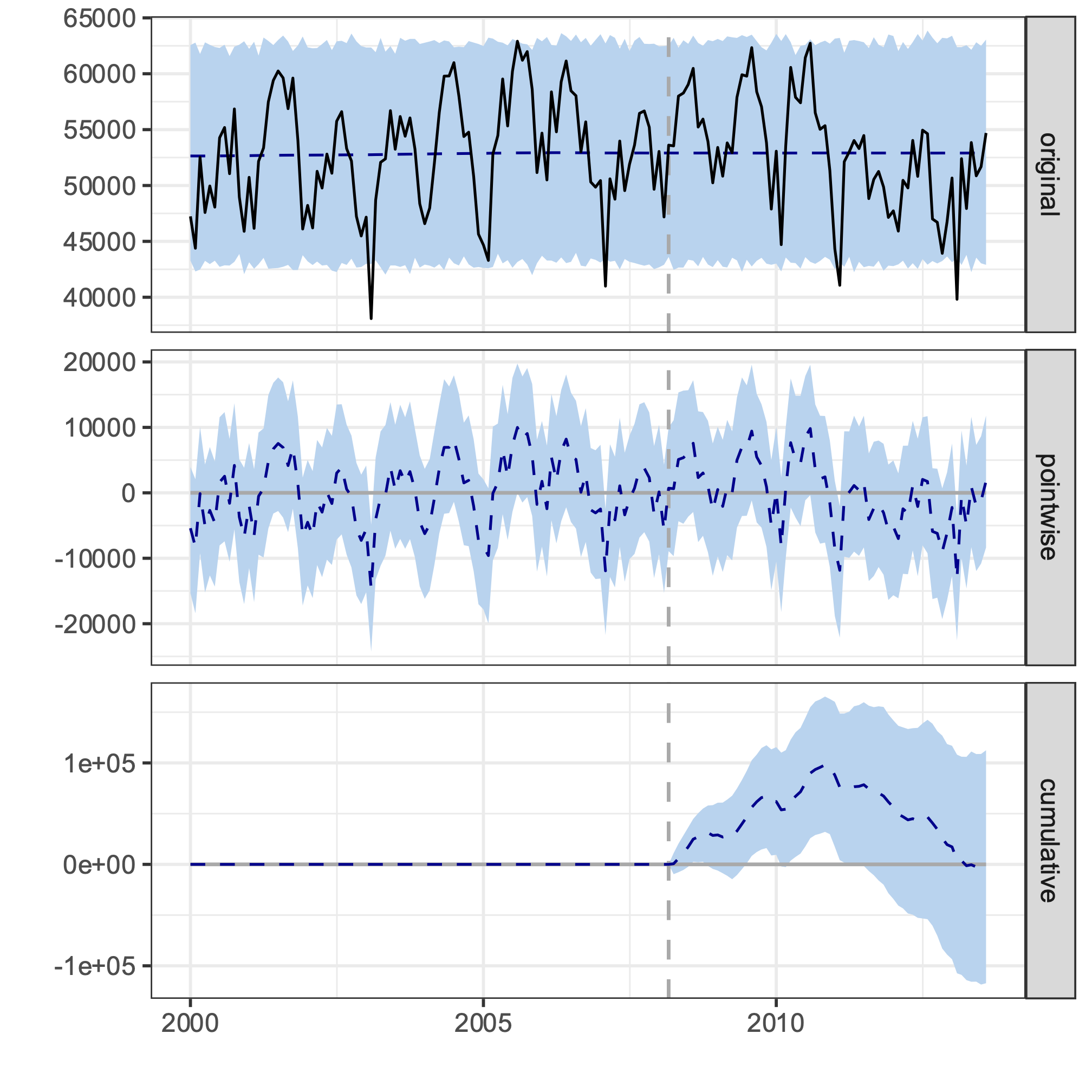

Figure S 8 Determining causal impact on behavior data with a paired covariate

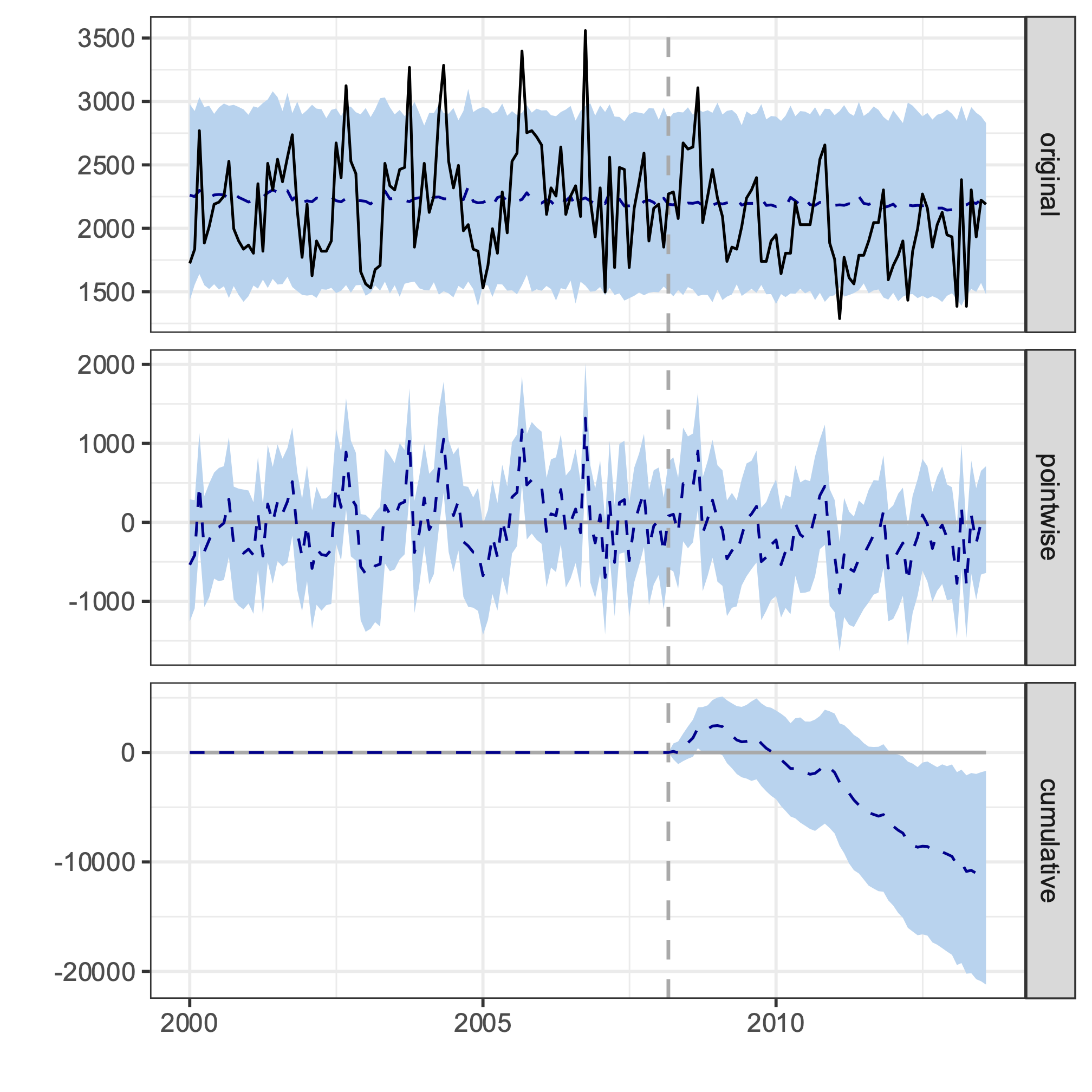

### *Section 4. Additional Information Regarding the Biomedical (Diabetes Data) Analysis*

Choosing specific covariates for the Bayesian Structural Time Series model of the clinical diabetes data involved aggregating variables by 24-hour intervals via statistical transformations. Below we show the correlation matrices used to deduce relevant covariates.

Figure S 9 Correlation Matrices For Daily Transformations of Variables Against Daily Percent-Glucose-In-Target-Range And Each Other.

| 24-Hour Variance | |  |  |  |  |  |  |  |  |
| --- | --- | --- | --- | --- | --- | --- | --- | --- | --- |
| Percent Glucose  In Target Range | IOB | | HRV | StepCount | Active  Energy  Burned | Basal  Energy  Burned | Distance  Walking  Running | HeartRate | FlightsClimbed |
| 1 | 0.35565104 | | 0.04682685 | -0.0317146 | 0.07622221 | -0.1577995 | -0.0341479 | -0.0769831 | 0.42961495 |
| 0.35565104 | 1 | | 0.28510196 | -0.0670641 | -0.017652 | 0.16758298 | -0.0613314 | 0.05993794 | 0.09954991 |
| 0.04682685 | 0.28510196 | | 1 | 0.02413229 | 0.09871911 | -0.1628811 | 0.01832187 | 0.01323634 | -0.0525743 |
| -0.0317146 | -0.0670641 | | 0.02413229 | 1 | 0.02191346 | 0.15676812 | 0.99750163 | -0.0091451 | 0.10293797 |
| 0.07622221 | -0.017652 | | 0.09871911 | 0.02191346 | 1 | -0.075615 | 0.01676697 | -0.0702123 | 0.11221684 |
| -0.1577995 | 0.16758298 | | -0.1628811 | 0.15676812 | -0.075615 | 1 | 0.16750575 | -0.095641 | 0.01377467 |
| -0.0341479 | -0.0613314 | | 0.01832187 | 0.99750163 | 0.01676697 | 0.16750575 | 1 | -0.0108691 | 0.10296383 |
| -0.0769831 | 0.05993794 | | 0.01323634 | -0.0091451 | -0.0702123 | -0.095641 | -0.0108691 | 1 | -0.0346771 |
| 0.42961495 | 0.09954991 | | -0.0525743 | 0.10293797 | 0.11221684 | 0.01377467 | 0.10296383 | -0.0346771 | 1 |
| 24-Hour Mean | |  |  |  |  |  |  |  |  |
| Percent Glucose  In Target Range | IOB | | HRV | StepCount | Active  Energy  Burned | Basal  Energy  Burned | Distance  Walking  Running | HeartRate | FlightsClimbed |
| 1 | 0.38907708 | | -0.1122026 | 0.01914113 | 0.17664033 | -0.2209548 | 0.03844289 | 0.13838201 | 0.24232686 |
| 0.38907708 | 1 | | 0.04064072 | 0.07567834 | 0.16898891 | -0.1027674 | 0.09793497 | -0.0239459 | 0.09521906 |
| -0.1122026 | 0.04064072 | | 1 | -0.0475394 | -0.0710143 | 0.03670135 | -0.1067997 | -0.2738451 | 0.07395699 |
| 0.01914113 | 0.07567834 | | -0.0475394 | 1 | 0.07542843 | -0.122134 | 0.96681413 | 0.13361936 | 0.01600925 |
| 0.17664033 | 0.16898891 | | -0.0710143 | 0.07542843 | 1 | 0.01660937 | 0.0630408 | 0.14582858 | 0.07642928 |
| -0.2209548 | -0.1027674 | | 0.03670135 | -0.122134 | 0.01660937 | 1 | -0.1169834 | -0.0735549 | 0.07713834 |
| 0.03844289 | 0.09793497 | | -0.1067997 | 0.96681413 | 0.0630408 | -0.1169834 | 1 | 0.15758832 | -0.0325702 |
| 0.13838201 | -0.0239459 | | -0.2738451 | 0.13361936 | 0.14582858 | -0.0735549 | 0.15758832 | 1 | 0.06692821 |
| 0.24232686 | 0.09521906 | | 0.07395699 | 0.01600925 | 0.07642928 | 0.07713834 | -0.0325702 | 0.06692821 | 1 |
| 24-Hour Minimum | |  |  |  |  |  |  |  |  |
| Percent Glucose  In Target Range | IOB | | HRV | StepCount | Active  Energy  Burned | Basal  Energy  Burned | Distance  Walking  Running | HeartRate | FlightsClimbed |
| 1 | -0.0561953 | | -0.1641335 | 0.12209808 | -0.0921188 | -0.0998087 | 0.16460823 | 0.13178686 | -0.0715939 |
| -0.0561953 | 1 | | 0.0736182 | 0.45421359 | 0.02248755 | 0.13963123 | 0.42173216 | 0.04546591 | -0.0167552 |
| -0.1641335 | 0.0736182 | | 1 | -0.0208383 | 0.15166274 | 0.14836172 | -0.0738506 | -0.0676928 | 0.0336735 |
| 0.12209808 | 0.45421359 | | -0.0208383 | 1 | 0.03279122 | 0.11140724 | 0.91775397 | -0.0048312 | 0.09669798 |
| -0.0921188 | 0.02248755 | | 0.15166274 | 0.03279122 | 1 | 0.53721547 | 0.00040976 | 0.28342752 | 0.10793276 |
| -0.0998087 | 0.13963123 | | 0.14836172 | 0.11140724 | 0.53721547 | 1 | 0.12237777 | 0.18942033 | -0.067886 |
| 0.16460823 | 0.42173216 | | -0.0738506 | 0.91775397 | 0.00040976 | 0.12237777 | 1 | -0.0367689 | -0.1318358 |
| 0.13178686 | 0.04546591 | | -0.0676928 | -0.0048312 | 0.28342752 | 0.18942033 | -0.0367689 | 1 | 0.16921989 |
| -0.0715939 | -0.0167552 | | 0.0336735 | 0.09669798 | 0.10793276 | -0.067886 | -0.1318358 | 0.16921989 | 1 |
| 24-Hour Maximum | |  |  |  |  |  |  |  |  |
| Percent Glucose  In Target Range | IOB | | HRV | StepCount | Active  Energy  Burned | Basal  Energy  Burned | Distance  Walking  Running | HeartRate | FlightsClimbed |
| 1 | 0.30403569 | | -0.01 | 0.00205067 | 0.06307793 | -0.234382 | 0.01486123 | 0.0297352 | 0.25353234 |
| 0.30403569 | 1 | | 0.09899728 | -0.0903845 | 0.07541161 | -0.0080352 | -0.0884466 | 0.07258076 | 0.06006603 |
| -0.01 | 0.09899728 | | 1 | 0.01428585 | 0.13076999 | -0.126969 | -0.0291854 | 0.05033678 | -0.0438407 |
| 0.00205067 | -0.0903845 | | 0.01428585 | 1 | -0.0234174 | -0.0403758 | 0.97333865 | -0.035065 | 0.22202879 |
| 0.06307793 | 0.07541161 | | 0.13076999 | -0.0234174 | 1 | -0.1336745 | -0.0138854 | 0.11896388 | -0.0154114 |
| -0.234382 | -0.0080352 | | -0.126969 | -0.0403758 | -0.1336745 | 1 | -0.02301 | -0.0797334 | 0.06787426 |
| 0.01486123 | -0.0884466 | | -0.0291854 | 0.97333865 | -0.0138854 | -0.02301 | 1 | -0.0344696 | 0.18667004 |
| 0.0297352 | 0.07258076 | | 0.05033678 | -0.035065 | 0.11896388 | -0.0797334 | -0.0344696 | 1 | -0.0604643 |
| 0.25353234 | 0.06006603 | | -0.0438407 | 0.22202879 | -0.0154114 | 0.06787426 | 0.18667004 | -0.0604643 | 1 |
| 24-Hour Range | |  |  |  |  |  |  |  |  |
| Percent Glucose  In Target Range | IOB | | HRV | StepCount | Active  Energy  Burned | Basal  Energy  Burned | Distance  Walking  Running | HeartRate | FlightsClimbed |
| 1 | 0.2972533 | | 0.08848623 | -0.0170331 | 0.06724451 | -0.17333 | -0.0117119 | -0.0152009 | 0.28383585 |
| 0.2972533 | 1 | | 0.12939748 | -0.0833513 | 0.07350533 | 0.14127527 | -0.0718342 | 0.0708919 | 0.03010398 |
| 0.08848623 | 0.12939748 | | 1 | 0.04711339 | 0.3150142 | -0.1737611 | 0.01961484 | 0.15844287 | -0.0597354 |
| -0.0170331 | -0.0833513 | | 0.04711339 | 1 | -0.010073 | 0.11763463 | 0.97820877 | -0.038455 | 0.23329806 |
| 0.06724451 | 0.07350533 | | 0.3150142 | -0.010073 | 1 | -0.1230368 | -0.0121033 | 0.12391679 | 0.02778952 |
| -0.17333 | 0.14127527 | | -0.1737611 | 0.11763463 | -0.1230368 | 1 | 0.15148451 | 0.03407447 | 0.07867423 |
| -0.0117119 | -0.0718342 | | 0.01961484 | 0.97820877 | -0.0121033 | 0.15148451 | 1 | -0.0406672 | 0.21970883 |
| -0.0152009 | 0.0708919 | | 0.15844287 | -0.038455 | 0.12391679 | 0.03407447 | -0.0406672 | 1 | -0.0023 |
| 0.28383585 | 0.03010398 | | -0.0597354 | 0.23329806 | 0.02778952 | 0.07867423 | 0.21970883 | -0.0023 | 1 |
| 24-Hour Sum | |  |  |  |  |  |  |  |  |
| Percent Glucose  In Target Range | IOB | | HRV | StepCount | Active  Energy  Burned | Basal  Energy  Burned | Distance  Walking  Running | HeartRate | FlightsClimbed |
| 1 | 0.39042109 | | -0.1113394 | 0.01936508 | 0.17799746 | -0.220228 | 0.03866424 | 0.14092842 | 0.24369841 |
| 0.39042109 | 1 | | 0.04138945 | 0.07557067 | 0.16802726 | -0.1047994 | 0.09781276 | -0.0226393 | 0.09510437 |
| -0.1113394 | 0.04138945 | | 1 | -0.0473129 | -0.0702472 | 0.03654206 | -0.1065559 | -0.2643036 | 0.0759667 |
| 0.01936508 | 0.07557067 | | -0.0473129 | 1 | 0.07532169 | -0.1225852 | 0.96681058 | 0.13284762 | 0.0161177 |
| 0.17799746 | 0.16802726 | | -0.0702472 | 0.07532169 | 1 | 0.01456977 | 0.06289779 | 0.1451525 | 0.07619192 |
| -0.220228 | -0.1047994 | | 0.03654206 | -0.1225852 | 0.01456977 | 1 | -0.1174446 | -0.0760203 | 0.07553856 |
| 0.03866424 | 0.09781276 | | -0.1065559 | 0.96681058 | 0.06289779 | -0.1174446 | 1 | 0.15645926 | -0.0324614 |
| 0.14092842 | -0.0226393 | | -0.2643036 | 0.13284762 | 0.1451525 | -0.0760203 | 0.15645926 | 1 | 0.07266538 |
| 0.24369841 | 0.09510437 | | 0.0759667 | 0.0161177 | 0.07619192 | 0.07553856 | -0.0324614 | 0.07266538 | 1 |
